# Pandemic danger to the deep: the risk of marine mammals contracting SARS-CoV-2 from wastewater

**DOI:** 10.1101/2020.08.13.249904

**Authors:** Sabateeshan Mathavarajah, Amina K. Stoddart, Graham A. Gagnon, Graham Dellaire

**Author notes:** **Corresponding author:** Graham Dellaire, Ph.D., Departments of Pathology and Biochemistry & Molecular Biology, Dalhousie University, P.O. BOX 15000, Halifax, Nova Scotia, Canada, B3H 4R2.

## Abstract

We are in unprecedented times with the ongoing COVID-19 pandemic. The pandemic has impacted public health, the economy and our society on a global scale. In addition, the impacts of COVID-19 permeate into our environment and wildlife as well. Here, we discuss the essential role of wastewater treatment and management during these times. A consequence of poor wastewater management is the discharge of untreated wastewater carrying infectious SARS-CoV-2 into natural water systems that are home to marine mammals. Here, we predict the susceptibility of marine mammal species using a modelling approach. We identified that many species of whale, dolphin and seal, as well as otters, are predicted to be highly susceptible to infection by the SARS-CoV-2 virus. In addition, geo-mapping highlights how current wastewater management in Alaska may lead to susceptible marine mammal populations being exposed to the virus. Localities such as Cold Bay, Naknek, Dillingham and Palmer may require additional treatment of their wastewater to prevent virus spillover through sewage. Since over half of these susceptibility species are already at risk worldwide, the release of the virus via untreated wastewater could have devastating consequences for their already declining populations. For these reasons, we discuss approaches that can be taken by the public, policymakers and wastewater treatment facilities to reduce the risk of virus spillover in our natural water systems. Thus, we indicate the potential for reverse zoonotic transmission of COVID-19 and its impact on marine wildlife; impacts that can be mitigated with appropriate action to prevent further damage to these vulnerable populations.

## 1. Introduction

In many municipalities, sewage whether treated or untreated is the major effluent discharged into rivers and oceans after agricultural run-off. Although the “seaward” design of sanitation systems is ideal for our public health, the same is not true for ocean life. Marine mammals are vulnerable to the influx of pharmaceuticals, personal care products, detergents and nanoparticles from sewage. Another invisible threat carried in the sewage to marine life is the potential influx of foreign microorganisms found in human excrement. During the current pandemic, the foreign pathogen of interest is the Severe acute respiratory syndrome coronavirus 2 (SARS-CoV-2); a virus that originated in Wuhan, China in December 2019 and is responsible for coronavirus disease 2019 (COVID-19)(Yang et al., 2020).

It is well-established that the SARS-CoV-2 virus infects the respiratory system but recent evidence indicates that the virus is responsible for multi-organ infection, including the intestinal tract of patients, which results in gastroenteritis in one third of COVID-19 cases (Lamers et al., 2020). The viral shedding in feces occurs irrespective of symptoms and age. A consequence of its capacity to infect the human gut and the shedding is the presence of viable coronavirus in the human waste of infected individuals, and thus a potential source of viral transmission (Fei Xiao et al., 2020). Indeed, it has been show that COVID-19 patients shed infective SARS-CoV-2 in stool and urine (Sun et al., 2020; Fei Xiao et al., 2020). The importance of this potential mechanisms of transmission is further highlighted by the fact that SARS-CoV-2 has been detected widely in the untreated domestic wastewater of many countries including Spain, Australia, Italy and France (Ahmed et al., 2020; La Rosa et al., 2020; Randazzo et al., 2020).

Domestic wastewater is treated through primary, secondary and tertiary treatment stages. Primary treatment involves the settling of settleable solids and is used alone or in conjunction with subsequent secondary and tertiary treatment stages. As such, primary treatment alone is predicted to poorly mitigate viral contamination. During secondary treatment, a biological process is used to remove dissolved and suspended organic compounds. During tertiary treatment, various processes are used to further reduce nutrient and pathogen concentrations. SARS-CoV-2 RNA can be easily detected in treated wastewater effluent, and initial data suggests that treatment stage does impact SARS-CoV-2 RNA, which in turn may serve as a measure of viral load (Randazzo et al., 2020). For example, using primary sludge to detect initial levels of SARS-CoV-2 in environmental systems, one study found that 83% (35 out of 45) of primary samples tested positive for SARS-CoV-2 RNA, 11% (2 out of 18) secondary treated effluent samples tested positive for SARS-CoV-2, while none of the tertiary treated effluent samples (0 out of 12) tested positive (Peccia et al., 2020). In adapting to the pandemic, there is a need to identify wastewater treatment plants that are at relatively high risk of allowing effluent carrying infective SARS-CoV-2 to “spillover” or “overflow” into natural systems, as we show in our study. Thus, non-treated and primary-treated wastewater represents the highest risk effluent for SARS-CoV-2 transmission.

SARS-CoV-2 can be detected in our sewage and efforts are being made to utilize wastewater virus detection to manage responses to COVID-19 and potentially future pandemics. For example, in Italy, SARS-CoV-2 was recently detected in untreated wastewater, and as such, its survival in water and wastewater represents a significant form of transmission to consider (La Rosa et al., 2020). Furthermore, in Paris it was shown that high concentrations of SARS-CoV-2 RNA in sewage water between March and April 2019 correlated with a spike in deaths from COVID-19 approximately 7 days later, suggesting that the detection of virus in wastewater represents a possible early warning system for COVID-19 and future viral pandemics (Wurtzer et al., 2020). On June 16th, it was reported that SARS-CoV-2 was detected in the river water in Ecuador (a country with poor sanitation), where the untreated sewage is delivered directly into natural waters (Guerrero-Latorre et al., 2020). Although the situation in Ecuador is one example, the challenges of managing direct sewage outfalls during the pandemic in sub-Saharan Africa highlight the global calls to action of understand SARS CoV-2 in natural water systems (Street, Malema, Mahlangeni, & Mathee, 2020).

The obvious question arises from these studies, does SARS-CoV-2 viral RNA load equate to infective virus and thus risk? Previously, it was shown that other related coronaviruses are stable in wastewater and water from days to weeks (Casanova, Rutala, Weber, & Sobsey, 2009). A study found that the SARS-CoV-2 virus is stable in a wide range of pH (3-10), including water (A. W. H. Chin et al., 2020). However, how the SARS-CoV-2 virus fares in the environment and the conditions influencing its stability in wastewater are still poorly understood. Casanova et al., 2009 found that in lake water, another coronavirus, mouse hepatitis virus (MHV), was stable for longer than 2 weeks. Finally, modelling studies by Shutler and colleagues that extrapolate infectivity from *in vitro* studies suggest that SARS-CoV-2 viral particle may be stable for up to ~25 days in warmer regions of the world (Shutler et al., 2020). The study also predicted how differing environments around the world differ in relative risk of the virus in wastewater discharge (Shutler et al., 2020).

The sewage carrying infectious particles ends up in our natural water systems either directly discharged in some cases from inadequate treatment or from problematic sewage overflow or pipe exfiltration. Thus, the absence or failure of a wastewater treatment plant can lead to sewage being another form of transmission that affects both humans and susceptible species. Communities that do not have a safe sanitation infrastructure in place will put many individuals and susceptible species at risk. A recent preprint provided the first evidence of SARS-CoV-2 transmission through sewage, which lead to the infection of multiple individuals in a low-income community in Guangzhou, China (Yuan et al., 2020). SARS-CoV-2 was found in street sewage puddles and pipes in the area. In a similar way, infectious particles from the sewage ending up in our natural waters may lead to virus spillover for susceptible species. Collectively, these studies indicate the potential for SARS-CoV-2 to be stable enough in wastewater for raw sewage to be a contributor to virus transmission during the pandemic.

Historically, wastewater management in the context of infectious disease has been confined to local infrastructures. Historically relevant waterborne infectious species such as *Escherichia coli, Giardia lamblia* and *Cryptosporidium spp*. are assessed locally, and the appropriate actions/treatments can be taken or utilized (Mayer & Palmer, 1996; Nasser, Vaizel-Ohayon, Aharoni, & Revhun, 2012). However, in the case of an infectious virus with a global reach being present in our wastewater, we are in a new paradigm. Like the virus, we must continue to globally evolve wastewater management practices and risk assessment to protect not only us but other organisms that are predicted to be susceptible to the virus. Infectious SARS-CoV-2 virus has been isolated from both urine and fecal samples; new indicators that the virus is being shed into sewage (Sun et al., 2020; F. Xiao et al., 2020). Consequently, it is possible that marine mammal species that are found inland or nearshore of contaminated natural water systems will be exposed to a novel pandemic virus. But what is the likelihood of such a scenario?

A potential spillover of a human pandemic-type virus to wild animals via reverse zoonosis can have drastic consequences on their populations. A recent example occurred with the Ebola pandemic where over 5000 gorillas were lost over the course of 2 years (2002 and 2003) after a human strain of the virus was transmitted to a gorilla (Bermejo et al., 2006; Wolinsky, 2017). There is a need to prevent such spillover events from occurring to protect vulnerable species around the world and that begins with assessing the possibility of it occurring for species in proximity to humans. The transmission of the SARS-CoV-2 virus from humans to other vulnerable species in proximity to humans has already been observed. In a New York Public zoo, a critically endangered tiger and several other animals (including lions) were infected with the SARS-CoV-2 virus (Wang, Mitchell, et al., 2020). When the perpetrator virus was sequenced from a sick Malayan tiger, it was revealed that it was two strains of human origin (i.e, two separate human-feline transmission events) that had infected these feline species (McAloose et al., 2020). Furthermore, there is now evidence that the virus can infect and be carried by certain domesticated pets, including cats; felines that are closely related to tigers (Shi et al., 2020). We recently described how ACE2 (the host receptor targeted by SARS-CoV-2) variability has contributed to why certain species are susceptible and not others (Mathavarajah & Dellaire, 2020). We then used these differences to predict the susceptibility of other at-risk feline species (Mathavarajah & Dellaire, 2020). In addition, a report utilizing a similar approach suggested that only mammals were predicted to be highly susceptible to SARS-CoV-2 and at risk (Damas et al., 2020). In support of these analyses, chickens were recently found to not be susceptible to the virus (Schlottau et al., 2020). Therefore, with the recent studies on wastewater and the presence of infective SARS-CoV-2, it is increasingly likely that wildlife, and in particular mammals exposed to waterways receiving wastewater effluent, will be exposed to this virus. As such, additional public efforts to prevent spillover events will be beneficial for protecting susceptible marine species during the current COVID-19 pandemic, particularly those species that are threatened or endangered. Thus, it is important to identify which marine mammals are at risk of contracting the virus. In addition, identifying localities with non-treated and primary-treated wastewater effluent and their proximity to endangered marine mammal populations will be important for risk-mitigation. In this article, we explore the susceptibility of marine mammal species, the conservation risks associated with the anthropogenic transmission of the SARS-CoV-2 virus to marine mammals around the world, and examine Alaskan wastewater management practices as a case study for identifying localities with a potentially high risk of virus spillover.

## 2. Methods

### 2.1 ACE2 variation in marine mammals

We performed a comprehensive analysis of the ACE2 receptor using annotated genomes of marine mammals from the 4 major groups: *Cetacea* (whales and dolphins), *Pinnepidia* (seals), *Sirenia* (sea cows), and *Fissipedia* (sea otters and polar bears). In total, we examined 36 marine mammal species that encompassed all publicly available reference and scaffold genomes. We compared the predicted ACE2 orthologs from these genomes to identify whether or not binding residues (25 consensus residues) are mutated (Shang et al., 2020). The analysis extends previous work by Damas et al., (2020) where they looked at the ACE2 receptor in 12 cetacean species. Our goal with these analyses was to generate the most comprehensive index of susceptibility for marine mammals to SARS-CoV-2 based on *ACE2* gene conservation.

### 2.2. Assessing the susceptibility of species to SARS-CoV-2

After compiling the mutation data for comparisons between human ACE2 and the orthologous ACE2 sequences, we utilized the MutaBind2 tool to model and predict whether these mutations (i) improve or (ii) reduce SARS-CoV-2 binding affinity to the receptor (Mihindukulasuriya, Wu, St Leger, Nordhausen, & Wang, 2008; Zhang et al., 2020). The reference protein for the analysis is the human ACE2 receptor. MutaBind2 models how protein-protein interactions are affected from missense mutations in the protein sequence. The RBD binding domain of the SARS-CoV-2 virus spike protein interacts with the human ACE2 receptor to facilitate virus entry. For that reason, we used the crystal structure of the Spike protein and ACE2 receptor as a template for the modelling (PDB: 6M17) (Yan et al., 2020). If the ACE2 orthologs carry mutations that reduce the binding affinity of the Spike protein to ACE2, then we predict susceptibility to be reduced.

To distinguish between the susceptible and non-susceptible species, we used the human ACE2 (high susceptibility), feline ACE2 (medium susceptibility; lower affinity but still susceptible) and dog ACE2 (not susceptible) as reference points to stratify groups in terms of susceptibility based on ΔΔGbind (kcal mol^−1^) values. Here, ΔΔGbind describes the predicted change in binding affinity of the SARS-CoV-2 Spike protein induced by mutations in the ACE2 receptor. Susceptibility values based on ΔΔGbind were determined as follows: higher than human (<−0.1 kcal mol^−1^), high (resembles human ACE2; −0.1 to 0.1 kcal mol^−1^), medium (resembles cat ACE2; 0.1 to 0.4 kcal mol^−1^), lowly (resembles dog ACE2; >0.4 kcal mol^−1^). Negative values are stabilizing, and positive values are destabilizing mutations on the proteinprotein interaction. Although these are relative values, they allow us to compare susceptibilities of species based on available data. The analysis thus allows the prediction of which species have high or low susceptibility to virus entry via ACE2 and ultimately an estimate of the degree a species is at-risk. These data and analyses are summarized in Supplemental Table 1.

### 2.3. Cross-referencing of conservation status and susceptibility

We cross-referenced the International Union for Conservation of Nature (IUCN) Red list of Threatened Species (https://www.iucnredlist.org; accessed July 31, 2020) to better understand which at risk species were also susceptible to the virus. For each species, we looked at the global assessment to determine their IUCN Red List Category. Only Near Threatened, Vulnerable, Endangered and Critically Endangered species were included for comparison. Some species did not have enough global data for assessment (data deficient); these species are the *Orcinus orca* (Killer whale), *Kogia breviceps* (Pygmy sperm whale) and *Mesoplodon bidens* (Sowerby’s beaked whale). Moreover, it is important to note that these assessments of risk provide a worldwide overview of which species are at risk. Local populations need to be assessed because they may still be at risk from a potential virus spillover for susceptible species.

### 2.4. Geo-mapping of Alaska wastewater plants and species at risk

To identify high-risk areas for virus spillover in Alaska, we overlaid geo-mapping data with marine mammal population data. Wastewater treatment plant outfall location, treatment type, maximum daily flow, and name of receiving water body was assembled through permit search of the Alaska permit database maintained through the Alaska Department of Environmental and Conservation. These data are summarized in Supplemental Table 2. Two types of general National Pollutant Discharge Elimination System (NPDES) permits were used: i) small publicly owned treatment works and other small treatment works providing secondary treatment of domestic discharge to surface water (general permit number: AKG572000) and ii) lagoons discharging to surface water (general permit number: AKG57300). Wastewater treatment plants discharging through federally granted NPDES permits were not included in the dataset. The data on the marine mammal populations was derived from a 2016 study that observed 905 marine mammals and identified 13 different species in Alaska (Lefebvre et al., 2016). Only species analyzed in terms of susceptibility were included from the study’s population data in the analysis: (A) Northern fur seal (B) Stellar sea lions, (C) Northern sea otters, (D) Harbor seals, (E) Beluga whale, (F) Harbor porpoises, (G) Humpback whale.

## 3. Results

### 3.1. ACE2 variation in marine mammal species

Since marine mammals are at risk from wastewater carrying the virus, we sought out to comprehensively characterize their potential susceptibility to SARS-CoV-2 by examining all the publicly available sequenced genomes of marine mammal species. Our goal was to generate a thorough index identifying which marine mammal species are most at risk from waterborne athropogenic transmission of SARS-CoV-2, which could serve to urge additional cautionary measures for the protection of the vulnerable species. In all examined marine mammal genomes, both ACE2 binding residues D30 and M82 were mutated except for *Balaenoptera bonaerensis* (Anarctic Minke whale), which only has a M82 mutation (Fig. 1). However, these two mutations did not have a large destabilizing effect on the binding affinity of the SARS-CoV-2 spike protein to ACE2. The most mutations observed in the ACE2 consensus binding residues were identified in the *Zalophus californianus* (California Sea Lion) ACE2 receptor (8/25), which had a drastic reduction in virus binding affinity to the receptor (Fig.1). Similarly, *Sirenia* (sea cows) which only had one representative genome available, *Trichechus manatus* (West Indian Manatee), was predicted to have low susceptibility (7/25 residues mutated) (Fig.1).

**Figure 1.**
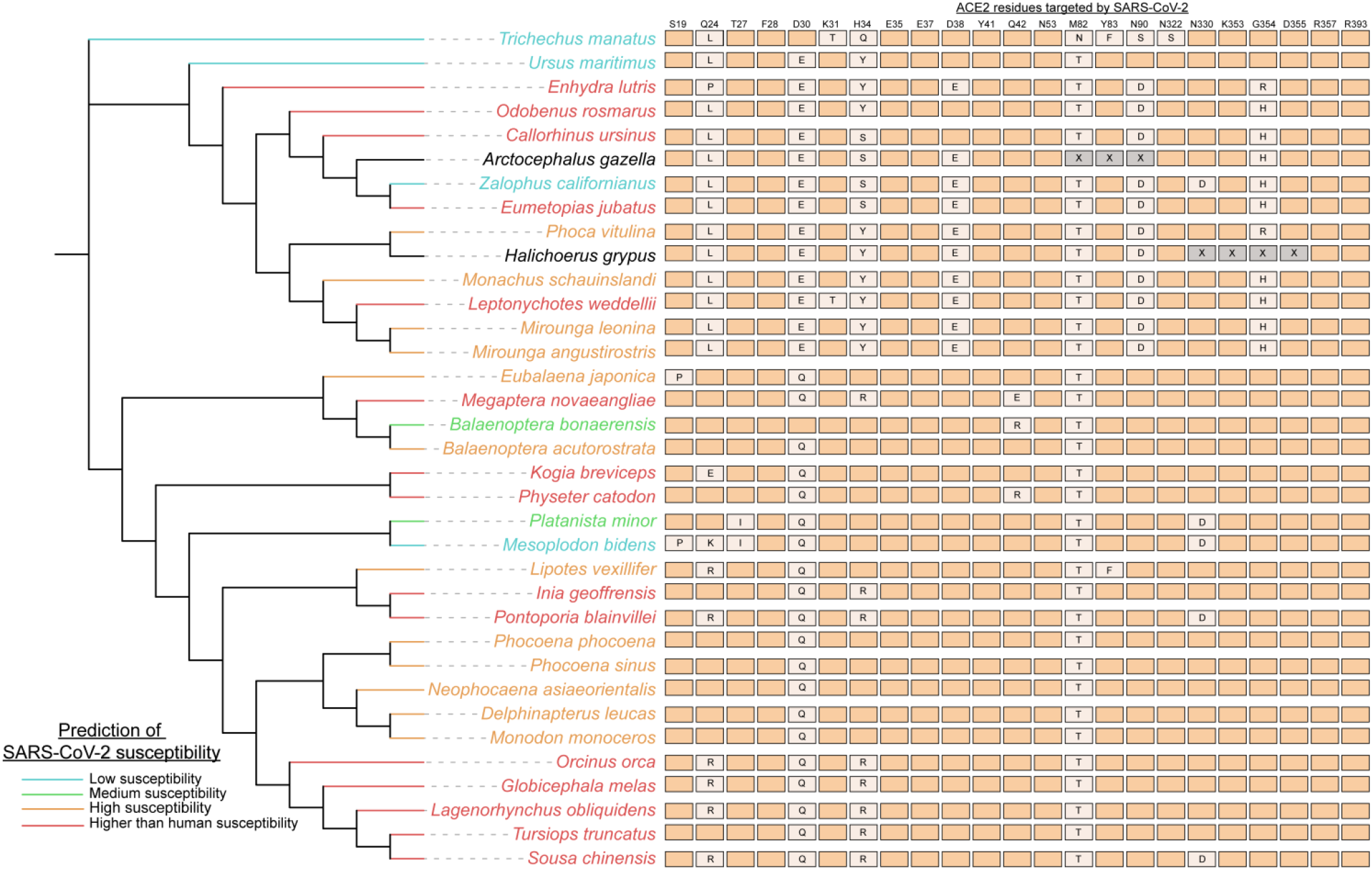
Marine mammal species are predicted to be susceptible to SARS-CoV-2. Species of Cetaceans, Fissipeds and Pinnipeds are predicted to be highly susceptible to the SARS-CoV-2 virus based on their ACE2 receptor. Genomes of 36 marine mammal species (all publicly available genomes for marine mammals on NCBI) were utilized to identify mutations in the ACE2 residues that bind SARS-CoV-2 to then determine susceptibility. The 20 consensus binding residues targeted by the SARS-CoV-2 virus are shown, with mutations denoted in orthologs noted (X= unknown residue; aligning regions were not available for *Arctocephalus gazelle* and *Halichoerus grypus* whole shotgun genome sequences). The susceptibility of species was scored using the MutaBind2 tool that determined how the mutations in ACE2 orthologs affected the binding affinity of the virus. Species were grouped in terms of susceptibility as follows: higher than human susceptibility (if binding affinity was increased by the mutations), high, medium and low. The phylogenetic tree and relationships between species were gathered from TimeTree (http://www.timetree.org) a database providing insight into the evolutionary relationships between species. Branch lengths are not representative of evolutionary times for the phylogenetic tree.

In contrast, there were a number of species (e.g, *Leptonychotes weddellii* (Weddell seal) and *Enhydra lutris* (sea otter)) with a relatively high number of binding residue mutations (6-8) but were predicted to have higher affinity binding to the SARS-CoV-2 spike protein by MutaBind2 (Fig.1), thus being potentially more susceptible to the SARS-CoV-2 virus that domestic cats used as our reference species. There are 15 species (of 36 examined) that have ACE2 receptors carrying mutations that further stabilize the interaction between the SARS-CoV-2 Spike protein and the ACE2 receptor. It is possible that these results reflect the origins of the virus in more closely related-mammals in evolutionary terms, likely bats and/or pangolins as reservoirs, which are more closely related to marine mammals than humans (Liu et al., 2020). Overall, we predict these 15 high-risk species that harbour stabilizing mutations in the ACE2 receptor will be the most vulnerable to SARS-CoV-2 infection if exposed.

### 3.2. *Cetacea, Pinnepedia* and *Fissipedia* susceptibility

Our analysis has predicted which species are most susceptible to the SARS-CoV-2 virus. The majority of *Cetacea* species (18/21 species) are predicted to have high or higher than human susceptibility to the virus (Fig. 1). ACE2 orthologs of cetaceans more so resemble the human ACE2 receptor in comparison to the other groups (Fig.1). Since many cetacean species are social, such as the *Tursiops truncates* (bottlenose dolphin) and *Delphinapterus leucas* (beluga whale), their high susceptibility suggests that their populations are especially vulnerable to intraspecies transmission of a novel virus such as SARS-CoV-2 (Fig.1). The *Mesoplodon bidens* (Sowerby’s beaked whale) was the only cetacean with predicted low susceptibility and has unique mutations in its ACE2 receptor such as S19P and Q24K (Fig.1). A group of closely related cetaceans (*Orcinus orca, Globicephala melas, Lagenorhynchus obliquidens, Tursiops truncatus*, and *Sousa chinensis*) are all predicted to more highly susceptible to the virus than humans. Therefore, many cetaceans appear to be at risk of contracting the virus if exposed.

Like *Cetacea*, the majority of *Pinnepedia* (seal) species (8/9) are also predicted to be highly susceptible to SARS-CoV-2 (Fig. 1). Two seal species, *Arctocephalus gazella* and *Halichoerus grypus*, had gaps in their orthologous *ACE2* gene and were not included in the analysis. However, many of their mutations resemble that of the other seal species that were predicted to be highly susceptible. The one exception is the lowly susceptible California sea lion, which has a novel N330D mutation in its orthologous ACE2 protein that appears to interfere with virus binding (Fig.1). It is also noteworthy that of all the examined marine mammal species, the *Odobenus rosmarus* (Atlantic Walrus) ACE2 is predicted to interact and bind with the greatest affinity to the SARS-CoV-2 spike protein (Fig.1). Among *Fissipedia, Enhydra lutris* (sea otters) are highly susceptible to the virus but polar bears are expected to have reduced susceptibility (Fig.1). We analyzed two sea otter subspecies genomes, the Northern (*E.lutris kenyoni*) and Southern (*E.lutris nereis*) sea otters and identified that both species have the same ACE2 mutations and were predicted to be highly susceptible. Thus, seals, walruses and otters represent at-risk species that may be impacted by waterborne anthropogenic transmission of SARS-CoV-2 virus.

### 3.3. Over half of species predicted to be susceptible to SARS-CoV-2 are already species at risk globally

Species that are at risk because of different anthropogenic sources are already the focus of conservation biologists around the world. To assess both the current risk of extinction for these species and the potential for SARS-CoV-2 spillover into their global populations, we cross-referenced their conservation status with their predicted susceptibility (Fig.2). We identified 15 species that are already at risk globally that fall under the categories of Near threatened, Vulnerable, Endangered and Critically endangered. The 15 species are predicted to be medium to higher susceptibility to the SARS-CoV-2 virus than humans. Only 2 of the 15 species fall under the “Near threatened” status (*Balaenoptera bonaerensis* and *Eumetopias jubatus*) (Fig.2). Furthermore, both the *Lipotes vexillifer* (Baiji) and *Phocoena sinus* (Vaquita) are critically endangered species that are predicted to have high susceptibility to the SARS-CoV-2 virus (Fig.2). Special care and attention are needed with our interactions with these species and with wastewater management in localities that are in proximity to habitats where populations from any of these 15 species are found. If these organisms are infected by SARS-CoV-2, their already dwindling population numbers will be even more at risk.

**Figure 2.**
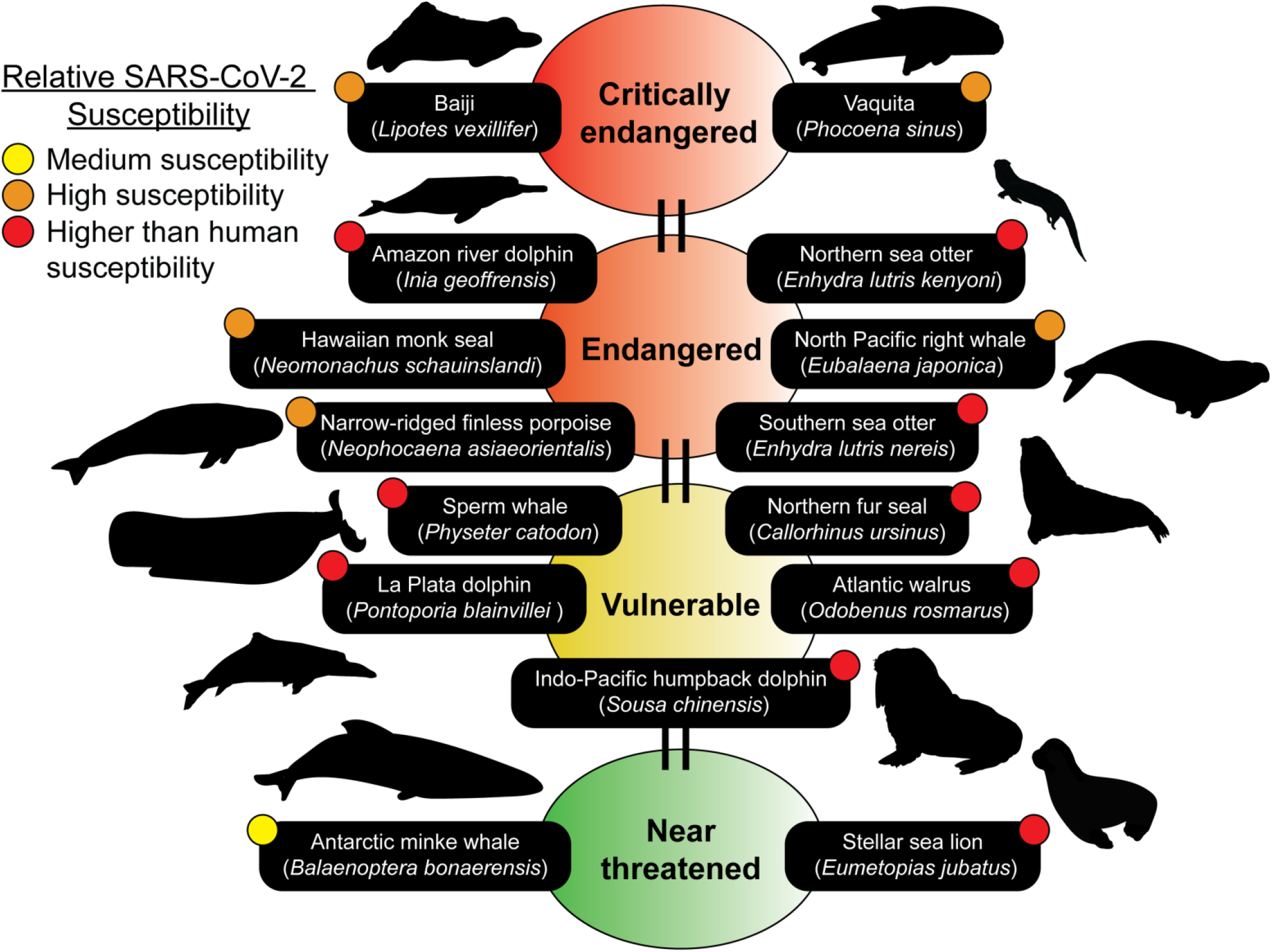
Marine mammal species predicted to be susceptible belong to the IUCN Red list. Many of the species predicted to be susceptible are members of the IUCN Red list of Threatened Species (https://www.iucnredlist.org). The IUCN Red list is an indicator of the world’s biodiversity and provides the most comprehensive data on the global conservation status of a species. 15 susceptible species ranging from medium to higher than human predicted susceptibilities can be identified on the IUCN Red list. Conservation statuses updated as of July 25^th^, 2020 were used. Silhouettes of species were drawn or obtained from PhyloPic (http://phylopic.org).

### 3.4. Wastewater treatment plants in Alaska and risk to endemic species

The identification of wastewater treatment plants and assessing the contribution of a municipality in terms of untreated wastewater discharge into natural water systems can help predict the potential hotspots for a spillover event. In Alaska, as of Aug 7^th^, 2020, there were 4221 confirmed cases of COVID-19 and this number continues to rise. Alaska is home to many marine mammal species, including a diverse variety of cetacean, seal and otter populations. Since there is a diversity of marine mammals in Alaska and their populations are well documented, we overlaid this information with available data on the wastewater treatment plants in this state. Using this approach, we were able to determine the potential geographic locations and species at high risk for anthropogenic transmission of SARS-CoV-2 via wastewater effluent in the state (Fig.3).

**Figure 3.**
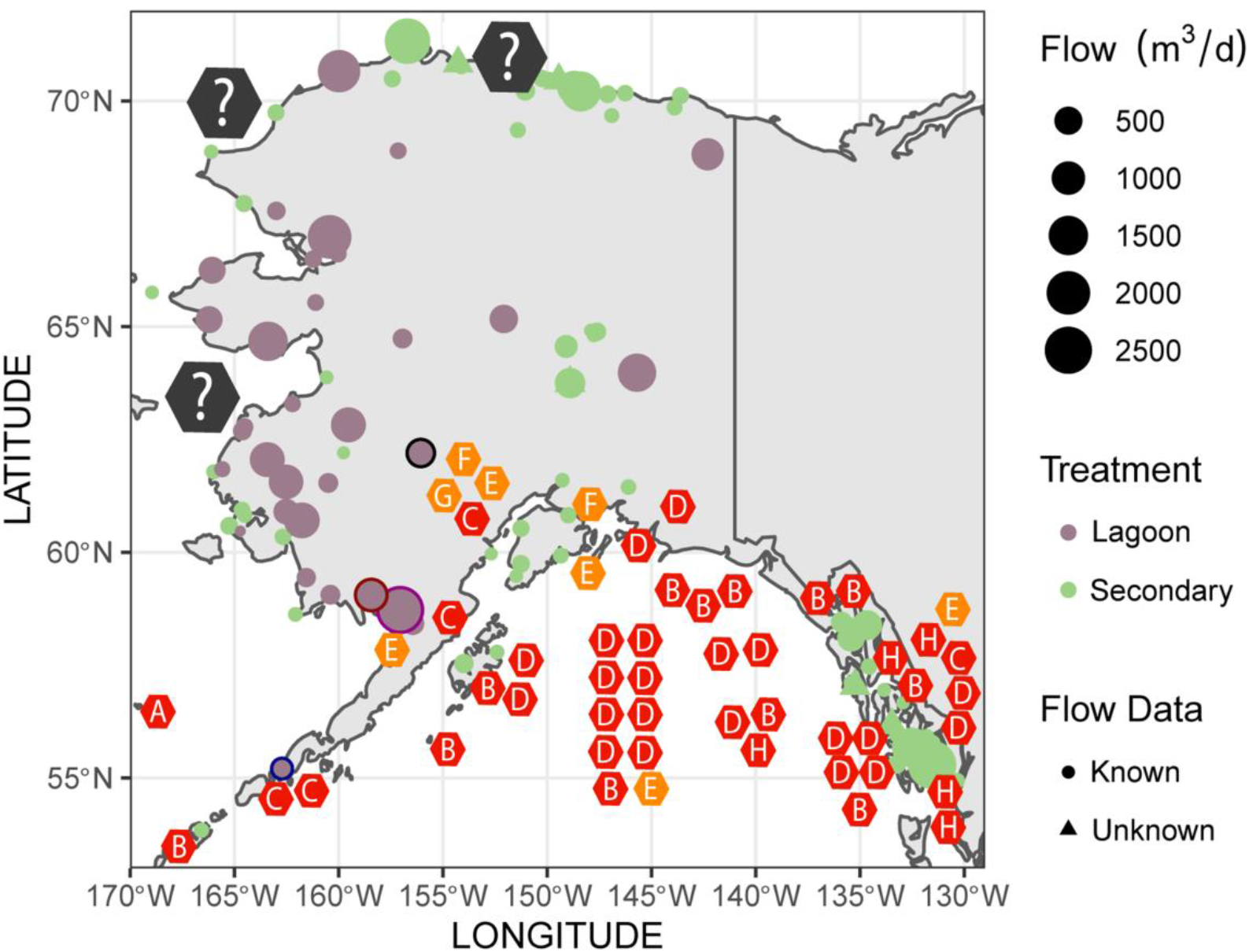
Wastewater management in Alaska identifies high risk areas for potential virus spillover. Geo-mapping of wastewater treatment plants was done using data from the Alaska Department of Environmental and Conservation. Types of treatment are listed for each plant (lagoon or secondary) and flow data is included. Overlap between species predicted to be susceptible and wastewater treatment plants that utilize lagoon treatment are identified as high-risk areas. High-risk sites and facilities discussed are bordered in black (Palmer; Talkeetna Lagoon), blue (City of Cold Bay; Cold Bay Lagoon), purple (Naknek; Naknek Lagoon) and red (Dillingham; Dillingham Lagoon). Marine mammal population data was obtained from a previous study (Lefebvre et al., 2016). Populations shown on the map in hexagons (represents 1-5 individuals observed at that location) and include the following species: (A) Northern fur seal (B) Stellar sea lions, (C) Northern sea otters, (D) Harbor seals, (E) Beluga whale, (F) Harbor porpoises, (G) Humpback whale. The colours of the hexagon species indicators describe predicted susceptibility; orange is high susceptibility and red is higher than human susceptibility. Grey hexagons indicate the *Phoca hispida* (Ringed seal), *Erignathus barbatus* (Bearded seal) and *Phoca largha* (Spotted seal) found in those areas. These species were not included in the analysis since their genomes are uncharacterized.

In Alaska, to treat wastewater there are lagoons (a form of primary treatment) and secondary treatment through wastewater treatment plants. The efficiency of lagoon treatment in dealing with SARS-CoV-2 has yet to be examined. However, SARS-CoV-2 has shown to be stable in a variety of environmental conditions and lagoon treatment is likely not sufficient for removing all infectious particles of the virus (Aboubakr, Sharafeldin, & Goyal, 2020; A. Chin et al., 2020). In contrast, secondary treatment appears to be efficient for removing most but does not remove all traces of SARS-CoV-2 RNA, as described Randazzo et al., (2020). We identified that municipalities of the northern and southern shores of Alaska rely mostly on secondary treatment for dealing with wastewater (Fig.3). In contrast, the western shores are dominated by lagoon treatment (Fig.3). In addition, the maximum effluent discharge flow rates are relatively high on the western shore, with many lagoons permitted to discharge from 2000-2500 m^3^/d of wastewater effluent (Fig.3). Three seal species are found along the western shore: the *Phoca hispida* (Ringed seal), *Erignathus barbatus* (Bearded seal) and *Phoca largha* (Spotted seal) (in order of prevalence). We are not able to predict the relative susceptibility of these species to SARS-CoV-2 due to an absence of publicly available genomic data for these species. However, one thing to note is that the Ringed seal and Spotted seal belong to the *Phoca* clade that are closely related to the harbor seal which was predicted to have high susceptibility to SARS-CoV-2 (Fig.1). Thus, the western shore has multiple high-risk areas for virus spillover, and it will be important going forward to gather data on their *ACE2* receptor sequence for the 3 seal species to properly assess potential risk in these populations.

Susceptible species examined in our study can be found primarily in the south shore of Alaska, within the Gulf of Alaska (Fig.3). Most of the wastewater treatment plants across the southern shores are secondary treatment plants that should efficiently remove SARS-CoV-2 (Fig.3). However, we have identified some high-risk locations in the south shore where lagoon treatment is used in the vicinity of marine mammals. These high-risk locations are the Cold Bay, Naknek, Dillingham and Palmer (Fig.3). The City of Cold Bay discharges wastewater into Cold Bay, where there are Northern sea otter populations (also found around the Aleutian Islands) that are predicted to be highly susceptible to the virus (Fig. 3). Similarly, beluga whales predicted to have high susceptibility can be found in Bristol Bay near Naknek, a city which relies only on lagoon treatment prior to the discharge of wastewater effluent into Bristol Bay (Fig.3). Like Naknek, Dillingham also discharges wastewater into Nushagak River where beluga whales are found (Fig.3) (Citta et al., 2016). In Palmer, wastewater effluent flows into the Talkeetna River, which is a tributary to the Susitna River and home to two species predicted to have high susceptibility, the beluga whales and harbor seals (Fig.3). The beluga whales found in Palmer belong to the Cook Inlet population that are currently endangered. Therefore, efforts to mitigate and carefully assess the impact of the wastewater effluent discharged only after primary treatment into theses marine mammal habitats will be important for protecting the species.

## 4. Discussion

Biological conservation focuses on assessing the impact of humans on biological diversity and determining how we can protect species that are at risk from extinction. Our impact on wildlife during a global pandemic may include the potential transmission of novel viruses to susceptible animals. In our study, we identified that half of the species predicted to be highly susceptible to SARS-CoV-2, based on ACE2 receptor conservation, are ones that are already at-risk species around the world. A potential reverse zoonoses event of SARS-CoV-2 may have grave implications for wildlife if not studied and mitigated. Currently there are many unknowns surrounding the biological effects and transmission of this novel virus in humans and other mammals. Thus, our study urges the need to act in a cautionary way during the current COVID-19 pandemic and future pandemics to protect marine wildlife through proper wastewater management.

Our analyses indicate that many different marine mammal species have an ACE2 receptor that is predicted to bind SARS-CoV-2 at a similar binding affinity as the human receptor, indicating that these species may contract the virus if encountered in their environment. The similarity in binding affinity stems from many of the key residues involved in the interaction being conserved. In addition, several of the ACE2 orthologs are predicted to bind tighter than the human ACE2 receptor with the SARS-CoV-2 virus (i.e, higher than human susceptibility) and it is possible that infection may occur at lower titres of the virus in these species. Based on the studies looking at how cetaceans and seals are affected by other coronaviruses, the pathological outcomes may be devastating for certain species. Infection with other coronaviruses in these species leads to severe necrosis of the liver (in beluga whales) and lung necrosis (in Pacific harbor seals) (Mihindukulasuriya et al., 2008; Woo et al., 2014). In humans, research is pointing towards the notion that COVID-19 is also a multi-organ disease, with many different organs such as the lung, heart, kidneys, liver, brain, blood vessels and colon, all being affected after virus infection (Robba, Battaglini, Pelosi, & Rocco, 2020). Multi-organ infection was also shown for the related SARS-CoV-1 virus (Gu et al., 2005). Thus, the impacts of COVID-19 will likely extend from humans to susceptible wildlife and include the loss of life for susceptible marine mammals that succumb to multi-organ pathology.

The likelihood of further cross-species transmission of SARS-CoV-2, once in a marine species population, is also something to consider in conservation efforts. As discussed above, novel coronaviruses have already been found in cetaceans, indicating that viral disease is an important factor in their continued survival. These viruses belong to the *Gammacoronavirus* family, a group related to the *Betacoronavirus* family which include SARS-like viruses. The novel coronaviruses were identified in both the aforementioned social bottlenose dolphin and beluga whale (Mihindukulasuriya et al., 2008; Wang, Maddox, et al., 2020; Woo et al., 2014). These cetacean coronaviruses are related, and the strains identified in bottlenose dolphin were rapidly evolving, indicating a potential cross-species transmission event from beluga whales to the dolphins. Cases of cross-species transmission from humans to marine mammals has already been observed. Sea otters were implicated with the H1N1 pandemic (2009). Northern sea otters were tested two years after the pandemic and ~70% of the tested individuals were serologically positive for the Influenza A(H1N1)pdm09 virus (Li et al., 2014). The chain of transmission is complex and it appears that the Northern sea otters were likely infected after being in proximity to *Mirounga angustirostris* (Elephant seals) that also were infected by Influenza A(H1N1)pdm09 virus (Goldstein et al., 2013). In the case of the Elephant seals, a suggested hypothesis behind their infection with a human pandemic virus was their exposure to human excrement discharged in the water by shipping vessels in the area (Goldstein et al., 2013). Treatment measures may be required for shipping vessels during the COVID-19 pandemic since SARS-CoV-2 is also found in human waste. Furthermore, the chain of transmission highlights how anthropogenic factors have the capacity to cause the cross-species spillover of a pandemic-type virus. Furthermore, the interspecific transmission of the novel coronaviruses in both cetaceans and otters indicates the epizootic potential for their social species. Thus, the predicted high susceptibility to SARS-CoV-2 indicates that cetaceans may be at risk from a potential spillover of the current pandemic and if so, large portions of social cetacean populations could be affected.

Most seals are coastal species and for this reason, of concern with the current pandemic and their high susceptibility. In the past, an epizootic pneumonia that killed 21 *Phoca vitulina richardsii* (Pacific harbor Seal) seals off the coast of California was linked to a novel seal coronavirus (Nollens, Wellehan, Archer, Lowenstine, & Gulland, 2010). The lungs isolated from these seals showed signs of necrotizing lymphocytic interstitial pneumonia with intra-lesional bacteria. To make matters worse, the coronavirus was persistent within their populations and could be identified again 8 years later (Ng et al., 2011). Therefore, exposure of the seals to sewage-derived human SARS-CoV-2 virus in coastal waters could have similar long-lasting consequences for their populations and exacerbated by the fact that these animals are highly social.

In our analysis of wastewater management practices in Alaska, we identified that most of the wastewater treatment plants in the vicinity of marine mammals were utilizing secondary treatment, which likely rules out the possibility of virus exposure in these areas. However, there were certain locations that bordered marine mammal populations that only relied on primary treatment, which we identify as high-risk locations such as Palmer, Dillingham and Naknek. In Palmer, the beluga whale is at risk of exposure to the SARS-CoV-2 virus from wastewater effluent (Fig.3). The beluga whale population residing in Susitna River belong to an endangered population in the Cook Inlet that the river flows into (Goetz, Montgomery, Ver Hoef, Hobbs, & Johnson, 2012). In addition, although Anchorage (the most populated city in Alaska) wastewater treatment plants hold permits for secondary treatment, as of 2019, the permit of John M. Asplund Wastewater Treatment of Anchorage was modified to be primary (Anchorage Water and Waste Utility, 2019). Many Cook Inlet beluga whales have been observed to swim close to the effluent of the wastewater treatment plant and these animals may be exposed to SARS-CoV-2 from current wastewater treatment practices in Anchorage. Thus, Palmer and Anchorage wastewater management are concerning for the Cook Inlet beluga during the COVID-19 pandemic, one of the National Oceanic and Atmospheric Administration (NOAA) Fisheries’ “Species in the spotlight” - an initiative to save the most highly at-risk marine species. As such, wastewater discharge was already noted as one of the threats to the Cook Inlet beluga whales. A potential virus spillover into the Cook Inlet population of highly social and susceptible beluga whales may have devastating consequences for the success of their population moving forward. Thus, additional measures of treating the water in these areas would help protect the species and reduce the probability of them being exposed to SARS-CoV-2.

## 5. Conclusions and future directions

In our study, we determined that many marine mammal species are predicted to be susceptible to SARS-CoV-2 based on conservation of the virus host receptor ACE2. We then revealed how current wastewater management approaches in certain Alaskan localities may not be sufficient for preventing waterborne exposure of nearby marine mammals to the virus. Our work urges for greater caution in considering how wastewater management practices can play a role in disease transmission, including reverse zoonotic transmission of waterborne pathogens from humans to susceptible non-human species during pandemics.

Given the grave consequences of waterborne anthropogenic transmission of SARS-CoV-2 to marine populations, are there concrete steps we can take to mitigate the risks to wild and captive marine animal populations from current and future pandemics? To begin to address this question, we have assembled a list of possible approaches (Table 1). For example, restricting contact and access to at-risk species in marine parks could help protect captive marine mammals. Such an approach has been suggested to protect other susceptible species including lions and tigers, following the reverse zoonotic infection of a tiger with SARS-CoV-2 at the Bronx Zoo in New York (Wang, Mitchell, et al., 2020). Similarly, non-human primate Old world apes and monkeys are also predicted to be highly susceptible to SARS-CoV-2 and these findings have led to improved measures for protecting these primates at zoos (Melin, Janiak, Marrone, Arora, & Higham, 2020). Such restriction measures should be present also in aquariums hosting susceptible marine mammals as well going forward.

**Table 1.** Approaches and research directions to consider going forward in the COVID-19 pandemic

| Future approaches | Target | Description |
| --- | --- | --- |
| SnotBot drones | Whales | SnotBot is a technology to study whale populations using a non-invasive approach (Wolinsky, 2017). Drones are utilized to collect whale snot to gather data on the genetics, microbiome and pregnancy status of the whale. The approach may also be useful for identifying infected whales. |
| Vaccination | Seals and other marine mammals | The vaccination approach was previously utilized in a feasible manner on Sable Island for wild grey seals, where it was used as a form of contraception (Brown et al., 1997). Since the technology is available, it may also be used for the purpose of protecting species from the COVID-19 pandemic. |
| Recovery of virus from wastewater | SARS-CoV-2 detection | Currently, the PEG, electronegative membrane, and ultrafiltration extraction methods are commonly utilized for the recovery of SARS-CoV-2 RNA from wastewater for qPCR (Warish Ahmed et al., 2020). These method ranges in recovery from 26.7 to 65.7% extraction of the related murine hepatitis virus RNA from the wastewater. These extraction methods still need to be assessed for recovery using SARS-CoV-2. Consequently, improving the methods for extracting both infective SARS-CoV-2 and viral RNA will allow for better estimation of the virus titres in wastewater. |
| Restrictions with captive animals | Marine mammals | Since many marine mammal species are predicted to be susceptible and a number of these species can be found in zoos and aquariums, there needs to be restrictions to limit contact between the public and animals. SARS-CoV-2 can be carried in aerosols and this may be a direct mechanism that leads to the infection of captive marine mammals by SARS-CoV-2. |
| Cell culture models | Marine mammals | Although we have predicted the susceptibility of marine mammal species to the SARS-CoV-2 virus, assessing their susceptibility using cell culture models may provide additional insight into the relative susceptibility of the species and also reveal how these species respond to the virus. |

For wild animal populations, the major concern will be the potential for wastewater exposure. We will need to improve our efforts towards monitoring and testing marine mammal populations to determine if any reverse zoonotic transmission events have occurred. New monitoring technologies will be powerful for such tasks. For example, SnotBot, where a drone retrieves mucus from whales, may be utilized to identify any virus spillover events since SARS-CoV-2 will likely be detected in the mucus of infected animals (Wolinsky, 2017). Non-invasive approaches like SnotBot are invaluable for the monitoring of virus spillover. Therefore, research into how we can continue monitoring these species post-pandemic will be paramount for assessing the impact of a human pandemic on wildlife.

We have described how potential high-risk spillover events can be identified using Alaska as a case study. If potential virus spillover is identified, a feasible approach may be vaccination of certain inland freshwater (e.g., otters) or coastal marine mammal populations (e.g., seals). The vaccination approach was previously shown to be feasible in a study of vaccination of Sable Island grey seals, where it was used as a form of contraception (Brown et al., 1997). Since the vaccination technology is available, it is not far-fetched to imagine that wild populations affected by the virus, just like humans, can be protected by herd immunity through vaccination.

Finally, the assessment and appropriate treatment of wastewater is a measure that will be key in reducing the effect of sewage transmission of the virus into natural water systems. Specifically, Shutler et al. (2020) outlines which countries pose the highest risk based on survival of the virus in wastewater. As such, high risk areas should act with caution and be attentive regarding how their wastewater is handled and treated. Similarly, we have described how analyzing localities (like Alaska) for susceptible species and their relative risk based on wastewater treatment can be helpful for determining localities of concern. In underdeveloped nations with poor sanitation, the major concern is on public health and the potential for the virus to contaminate human drinking water reservoirs. However, these nations are also particularly high risk for virus spillover if large amounts of raw sewage are discharged into natural water systems containing susceptible species. Thus, additional measures for treating sewage are important for both the health of the public and wildlife in these countries. Moreover, pinpointing locations of sewage overflows, and pipe leakage will allow for us to better assess where there is the most potential for wildlife to be affected. At this point in the pandemic, the available evidence indicates that wastewater is an important vector for SARS-CoV-2 transmission for humans and susceptible wildlife. Given the proximity of marine animals to high-risk environments where viral spill over is likely, we must act with foresight to protect marine mammal species predicted to be at-risk and mitigate the environmental impact of the COVID-19 pandemic.

## 6. Acknowledgments

We would like to thank Paul Wade (Marine Mammal Laboratory, NOAA) for his comments and helpful insight on the Cook Inlet beluga whales, as well as Jeremie Saunders and the SickBoy Podcast hosted by the Canadian Broadcasting Corporation (CBC) for drawing attention to this work and the risks of COVID-19 transmission through wastewater. This work is supported by a Discovery Grant and Alliance Grant from the Natural Sciences and Engineering Research Council of Canada (NSERC)(RGPIN-04034) to G.D. and (ALLRP 554503-20) to A.S and G.G., and S.M. is supported by a Killam Doctoral Award, as well as a Nova Scotia Graduate Scholarship and Dalhousie University’s Presidents Award.

## 7. Author Contributions

G.D. and S.M. conceived of the study; S.M performed the bioinformatics analysis; A.S performed the geo-mapping and characterized wastewater treatment plants; All authors wrote and edited the manuscript.

## Abbreviations

ACE2: Angiotensin converting enzyme 2
RBD: receptor binding domain
CoVID-19: Coronavirus Disease-2019

## References

Aboubakr, H. A., Sharafeldin, T. A., & Goyal, S. M. (2020). Stability of SARS-CoV-2 and other coronaviruses in the environment and on common touch surfaces and the influence of climatic conditions: A review. Transbound Emerg Dis. doi:10.1111/tbed.13707

Ahmed, W., Angel, N., Edson, J., Bibby, K., Bivins, A., O’Brien, J. W.,… Mueller, J. F. (2020). First confirmed detection of SARS-CoV-2 in untreated wastewater in Australia: A proof of concept for the wastewater surveillance of COVID-19 in the community. Sci Total Environ, 728, 138764. doi:10.1016/j.scitotenv.2020.138764

Anchorage Water and Waste Utility. (2019). 2019 Asplund Wastewater Treatment Facility Annual Monitoring Report. Retrieved from https://www.awwu.biz/water-quality/cook-inlet-water-quality

Bermejo, M., Rodriguez-Teijeiro, J. D., Illera, G., Barroso, A., Vila, C., & Walsh, P. D. (2006). Ebola outbreak killed 5000 gorillas. Science, 314(5805), 1564. doi:10.1126/science.1133105

Brown, R. G., Bowen, W. D., Eddington, J. D., Kimmins, W. C., Mezei, M., Parsons, J. L., & Pohajdak, B. (1997). Evidence for a long-lasting single administration contraceptive vaccine in wild grey seals. J Reprod Immunol, 35(1), 43–51. doi:10.1016/s0165-0378(97)00047-8

Casanova, L., Rutala, W. A., Weber, D. J., & Sobsey, M. D. (2009). Survival of surrogate coronaviruses in water. Water Res, 43(7), 1893–1898. doi:10.1016/j.watres.2009.02.002

Chin, A., Chu, J., Perera, M., Hui, K., Yen, H.-L., Chan, M.,… Poon, L. J. m. (2020). Stability of SARS-CoV-2 in different environmental conditions.

Chin, A. W. H., Chu, J. T. S., Perera, M. R. A., Hui, K. P. Y., Yen, H.-L., Chan, M. C. W.,… Poon, L. L. M. (2020). Stability of SARS-CoV-2 in different environmental conditions. The Lancet Microbe, 1(1), e10–e10. doi:10.1016/S2666-5247(20)30003-3

Citta, J. J., Quakenbush, L. T., Frost, K. J., Lowry, L., Hobbs, R. C., & Aderman, H. J. M. M. S. (2016). Movements of beluga whales (Delphinapterus leucas) in Bristol Bay, Alaska. 32(4), 1272–1298.

Damas, J., Hughes, G. M., Keough, K. C., Painter, C. A., Persky, N. S., Corbo, M.,… Lewin, H. A. (2020). Broad Host Range of SARS-CoV-2 Predicted by Comparative and Structural Analysis of ACE2 in Vertebrates. 2020.2004.2016.045302. doi:10.1101/2020.04.16.045302 %J bioRxiv

Goetz, K. T., Montgomery, R. A., Ver Hoef, J. M., Hobbs, R. C., & Johnson, D. S. J. E. S. R. (2012). Identifying essential summer habitat of the endangered beluga whale Delphinapterus leucas in Cook Inlet, Alaska. 16(2), 135–147.

Goldstein, T., Mena, I., Anthony, S. J., Medina, R., Robinson, P. W., Greig, D. J.,… Boyce, W. M. (2013). Pandemic H1N1 influenza isolated from free-ranging Northern Elephant Seals in 2010 off the central California coast. PLoS One, 8(5), e62259. doi:10.1371/journal.pone.0062259

Gu, J., Gong, E., Zhang, B., Zheng, J., Gao, Z., Zhong, Y.,… Leong, A. S. (2005). Multiple organ infection and the pathogenesis of SARS. J Exp Med, 202(3), 415–424. doi:10.1084/jem.20050828

Guerrero-Latorre, L., Ballesteros, I., Villacres, I., Granda-Albuja, M. G., Freire, B., & Rios-Touma, B. (2020). First SARS-CoV-2 detection in river water: implications in low sanitation countries. medRxiv, 2020.2006.2014.20131201. doi:10.1101/2020.06.14.20131201

La Rosa, G., Iaconelli, M., Mancini, P., Bonanno Ferraro, G., Veneri, C., Bonadonna, L.,… Suffredini, E. (2020). First detection of SARS-CoV-2 in untreated wastewaters in Italy. Sci Total Environ, 736, 139652. doi:10.1016/j.scitotenv.2020.139652

Lamers, M. M., Beumer, J., van der Vaart, J., Knoops, K., Puschhof, J., Breugem, T. I.,… Clevers, H. (2020). SARS-CoV-2 productively infects human gut enterocytes. Science. doi:10.1126/science.abc1669

Lefebvre, K. A., Quakenbush, L., Frame, E., Huntington, K. B., Sheffield, G., Stimmelmayr, R.,… Gill, V. (2016). Prevalence of algal toxins in Alaskan marine mammals foraging in a changing arctic and subarctic environment. Harmful Algae, 55, 13–24. doi:10.1016/j.hal.2016.01.007

Li, Z. N., Ip, H. S., Trost, J. F., White, C. L., Murray, M. J., Carney, P. J.,… Katz, J. M. (2014). Serologic evidence of influenza A(H1N1)pdm09 virus infection in northern sea otters. Emerg Infect Dis, 20(5), 915–917. doi:10.3201/eid2005.131890

Liu, Z., Xiao, X., Wei, X., Li, J., Yang, J., Tan, H.,… Liu, L. J. J. o. m. v. (2020). Composition and divergence of coronavirus spike proteins and host ACE2 receptors predict potential intermediate hosts of SARS-CoV-2. 92(6), 595–601.

Mathavarajah, S., & Dellaire, G. (2020). Lions, Tigers and Kittens too: ACE2 and susceptibility to CoVID-19. Evolution, Medicine, and Public Health, eoaa021. doi:10.1093/emph/eoaa021

Mayer, C. L., & Palmer, C. J. (1996). Evaluation of PCR, nested PCR, and fluorescent antibodies for detection of Giardia and Cryptosporidium species in wastewater. Appl Environ Microbiol, 62(6), 2081–2085.

McAloose, D., Laverack, M., Wang, L., Killian, M. L., Caserta, L. C., Yuan, F.,… Diel, D. G. (2020). From people to Panthera: Natural SARS-CoV-2 infection in tigers and lions at the Bronx Zoo. bioRxiv, 2020.2007.2022.213959. doi:10.1101/2020.07.22.213959

Melin, A. D., Janiak, M. C., Marrone, F., 3rd, Arora, P. S., & Higham, J. P. (2020). Comparative ACE2 variation and primate COVID-19 risk. bioRxiv. doi:10.1101/2020.04.09.034967

Mihindukulasuriya, K. A., Wu, G., St Leger, J., Nordhausen, R. W., & Wang, D. (2008). Identification of a novel coronavirus from a beluga whale by using a panviral microarray. J Virol, 82(10), 5084–5088. doi:10.1128/JVI.02722-07

Nasser, A. M., Vaizel-Ohayon, D., Aharoni, A., & Revhun, M. (2012). Prevalence and fate of Giardia cysts in wastewater treatment plants. J Appl Microbiol, 113(3), 477–484. doi:10.1111/j.1365-2672.2012.05335.x

Ng, T. F. F., Wheeler, E., Greig, D., Waltzek, T. B., Gulland, F., & Breitbart, M. (2011). Metagenomic identification of a novel anellovirus in Pacific harbor seal (Phoca vitulina richardsii) lung samples and its detection in samples from multiple years. J Gen Virol, 92(Pt 6), 1318–1323. doi:10.1099/vir.0.029678-0

Nollens, H. H., Wellehan, J. F., Archer, L., Lowenstine, L. J., & Gulland, F. M. (2010). Detection of a respiratory coronavirus from tissues archived during a pneumonia epizootic in free-ranging Pacific harbor seals Phoca vitulina richardsii. Dis Aquat Organ, 90(2), 113–120. doi:10.3354/dao02190

Peccia, J., Zulli, A., Brackney, D. E., Grubaugh, N. D., Kaplan, E. H., Casanovas-Massana, A.,… Omer, S. B. (2020). SARS-CoV-2 RNA concentrations in primary municipal sewage sludge as a leading indicator of COVID-19 outbreak dynamics. 2020.2005.2019.20105999. doi:10.1101/2020.05.19.20105999 %J medRxiv

Randazzo, W., Truchado, P., Cuevas-Ferrando, E., Simon, P., Allende, A., & Sanchez, G. (2020). SARS-CoV-2 RNA in wastewater anticipated COVID-19 occurrence in a low prevalence area. Water Res, 181, 115942. doi:10.1016/j.watres.2020.115942

Robba, C., Battaglini, D., Pelosi, P., & Rocco, P. R. M. (2020). Multiple organ dysfunction in SARS-CoV-2: MODS-CoV-2. Expert Rev Respir Med, 1–4. doi:10.1080/17476348.2020.1778470

Schlottau, K., Rissmann, M., Graaf, A., Schön, J., Sehl, J., Wylezich, C.,… Harder, T. J. T. L. M. (2020). SARS-CoV-2 in fruit bats, ferrets, pigs, and chickens: an experimental transmission study.

Shang, J., Ye, G., Shi, K., Wan, Y., Luo, C., Aihara, H.,… Li, F. (2020). Structural basis of receptor recognition by SARS-CoV-2. Nature, 581(7807), 221–224. doi:10.1038/s41586-020-2179-y

Shi, J., Wen, Z., Zhong, G., Yang, H., Wang, C., Huang, B.,… Bu, Z. (2020). Susceptibility of ferrets, cats, dogs, and other domesticated animals to SARS-coronavirus 2. Science. doi:10.1126/science.abb7015

Shutler, J., Zaraska, K., Holding, T. M., Machnik, M., Uppuluri, K., Ashton, I.,… Dahiya, R. (2020). Risk of SARS-CoV-2 infection from contaminated water systems. medRxiv, 2020.2006.2017.20133504. doi:10.1101/2020.06.17.20133504

Street, R., Malema, S., Mahlangeni, N., & Mathee, A. (2020). Wastewater surveillance for Covid-19: An African perspective. Science of The Total Environment, 743, 140719. doi:https://doi.org/10.1016/j.scitotenv.2020.140719

Sun, J., Zhu, A., Li, H., Zheng, K., Zhuang, Z., Chen, Z.,… Li, Y. M. (2020). Isolation of infectious SARS-CoV-2 from urine of a COVID-19 patient. Emerg Microbes Infect, 9(1), 991–993. doi:10.1080/22221751.2020.1760144

Wang, L., Maddox, C., Terio, K., Lanka, S., Fredrickson, R., Novick, B.,… Ross, K. (2020). Detection and Characterization of New Coronavirus in Bottlenose Dolphin, United States, 2019. Emerg Infect Dis, 26(7), 1610–1612. doi:10.3201/eid2607.200093

Wang, L., Mitchell, P. K., Calle, P. P., Bartlett, S. L., McAloose, D., Killian, M. L.,… Torchetti, M. K. (2020). Complete Genome Sequence of SARS-CoV-2 in a Tiger from a U.S. Zoological Collection. Microbiol Resour Announc, 9(22). doi:10.1128/MRA.00468-20

Wolinsky, H. (2017). Biology goes in the air: Unmanned aerial vehicles offer biologists an efficient tool for observation and sampling from a safe distance. EMBO Rep. doi:10.15252/embr.201744740

Woo, P. C., Lau, S. K., Lam, C. S., Tsang, A. K., Hui, S. W., Fan, R. Y.,… Yuen, K. Y. (2014). Discovery of a novel bottlenose dolphin coronavirus reveals a distinct species of marine mammal coronavirus in Gammacoronavirus. J Virol, 88(2), 1318–1331. doi:10.1128/JVI.02351-13

Wurtzer, S., Marechal, V., Mouchel, J.-M., Maday, Y., Teyssou, R., Richard, E.,… Moulin, L. (2020). Evaluation of lockdown impact on SARS-CoV-2 dynamics through viral genome quantification in Paris wastewaters. 2020.2004.2012.20062679. doi:10.1101/2020.04.12.20062679 %J medRxiv

Xiao, F., Sun, J., Xu, Y., Li, F., Huang, X., Li, H.,… Zhao, J. (2020). Infectious SARS-CoV-2 in Feces of Patient with Severe COVID-19. Emerg Infect Dis, 26(8). doi:10.3201/eid2608.200681

Xiao, F., Sun, J., Xu, Y., Li, F., Huang, X., Li, H.,… Zhao, J. J. E. i. d. (2020). Infectious SARS-CoV-2 in feces of patient with severe COVID-19. 26(8).

Yan, R., Zhang, Y., Li, Y., Xia, L., Guo, Y., & Zhou, Q. (2020). Structural basis for the recognition of SARS-CoV-2 by full-length human ACE2. Science, 367(6485), 1444–1448. doi:10.1126/science.abb2762

Yang, X., Yu, Y., Xu, J., Shu, H., Liu, H., Wu, Y.,… Yu, T. J. T. L. R. M. (2020). Clinical course and outcomes of critically ill patients with SARS-CoV-2 pneumonia in Wuhan, China: a single-centered, retrospective, observational study.

Yuan, J., Chen, Z., Gong, C., Liu, H., Li, B., Li, K.,… Liu, G. (2020). Coronavirus Disease 2019 Outbreak Likely Caused by Sewage Exposure in a Low-Income Community: Guangzhou, China, April 2020.

Zhang, N., Chen, Y., Lu, H., Zhao, F., Alvarez, R. V., Goncearenco, A.,… Li, M. (2020). MutaBind2: Predicting the Impacts of Single and Multiple Mutations on Protein-Protein Interactions. iScience, 23(3), 100939. doi:10.1016/j.isci.2020.100939

